# Phase Heterogeneity in Cholesterol-containing Ternary Phospholipid Lamellar Phases

**DOI:** 10.1101/2022.07.18.500534

**Authors:** Deborah L. Gater, Keontré I. Hughes, Vivian Stojanoff, A. F. Isakovic

## Abstract

Pseudo-ternary mixtures of lamellar phase phospholipids (DPPC and brain sphingomyelin with cholesterol) were studied below T_m_, while comparing the influence of cholesterol content, temperature and the presence of small quantities of Vitamin D binding protein (DBP) or Vitamin D receptor (VDR). The measurements, conducted by X-ray diffraction (XRD) and nuclear magnetic resonance (NMR), cover a range of cholesterol concentrations (20% mol. wt to 40% mol. wt.) and physiologically relevant temperature range (294 K – 314 K). In addition to rich intra-phase behaviour, data and modelling are used to approximate the lipids’ headgroup location variations under the above-mentioned experimental conditions.

## Introduction

Lamellar phase coexistence in ternary phospholipid systems containing cholesterol is a well-established phenomenon ^1^. In particular, the putative involvement of cholesterol and sphingomyelin in ‘lipid rafts’, or physiologically relevant domain formation/lateral lipid organisation *in vivo*, has resulted in a wealth of study in this area (see the following for more comprehensive background ^2,3^). However, much of this work has been conducted on fluid bilayers and less is understood about model lipid systems below their gel-fluid melting transition temperature (T_m_). The responses of ternary systems containing 1,2-dipalmitoyl-*sn*-glycero-3-phosphocholine (DPPC), sphingomyelin (SM) and cholesterol (Chol) ternary systems to neuroleptic drugs ^4^ and to phenothiazine derivatives ^5^ have been studied at lower Chol and SM concentrations than studied here. Evidence for microdomain formation, based on asymmetry in the phase transition observed by differential scanning calorimetry (DSC), was reported in large unilamellar vesicles (LUVs) containing DPPC/16:0-SM/Chol (85:10:5 mol%) ^4^. Similar evidence for phase separation was less clear in systems containing either 5% or 10% of both egg-yolk SM and Chol in DPPC, although the focus of that work was predominantly on inducing phase separation around the main DPPC transition temperature by the addition of phenothiazine derivatives ^5^. In both studies, the addition of small quantities (≤ 20 mol% total) of cholesterol and SM suppressed the DPPC pre-transition in DSC, and decreased the DPPC T_m_ (37.8 °C ^6^) by ∼ 1 °C.

Here, we present an analysis of x-ray diffraction (XRD) data obtained for multilamellar vesicles (MLVs) comprising different proportions DPPC, brain sphingomyelin (bSM) and Chol at three different temperatures between room temperature and physiological temperature (i.e. below or approaching the anticipated T_m_), with different combinations of buffer and added protein. The data are by nature complex for such systems, and thus we have sought to apply the methods of Harper *et al* ^7^ and of Rappolt ^8^ to estimate the range of structural parameters exhibited in these conditions. The XRD analysis is supported by ^31^P and ^13^C solid-state nuclear magnetic resonance (NMR) spectroscopy data.

The choice of added protein – Vitamin D receptor (VDR) or vitamin D binding protein (DBP) was motivated by a desire to commence a preliminary investigation of how lipid composition might affect the interactions of such proteins with phospholipid lamellar phases, and thus influence the transport and availability of vitamin D within the body. There have been numerous studies investigating the *in vitro* and *in vivo* relationships between, for example, vitamin D metabolism and phospholipid organisation^9^, or vitamin D bioavailability and factors including food/supplement matrix^10^. However, a detailed review of this area is beyond the scope of this article.

## Materials and Methods

### Materials

(DPPC), d_62_-DPPC and porcine bSM were bought from Avanti Polar Lipids (Alabama, US). Chol and D_2_O were purchased from Sigma Aldrich (US). Deionized water was used as solvent for all sample preparation unless otherwise specified. Vitamin D binding protein (DBP, Globulin GC) was bought from Athens Research (Georgia, US) & Vitamin D receptor (VDR) was from Abcam (UK). All materials were used without further purification.

### Sample Preparation

MLVs were formulated at desired mole percent ratios of DPPC, d_62_-DPPC, SM and Chol. MLVs were prepared by thin film hydration method where the blend of lipids was weighed out accurately and first dissolved together in a mixture of chloroform and methanol. The solvent was removed using a dry nitrogen stream and the samples were kept in vacuum overnight. The resulting thin film was then hydrated in deionized water. The hydrated samples were flash frozen using liquid nitrogen and lyophilized overnight. Dry samples were stored at -20 °C until needed. Samples were hydrated by the addition of H_2_O as indicated for characterization by XRD or NMR. Briefly, solvent in excess (66 wt%) was added directly to the lipids above 70 °C. After this, samples were heat-cycled twice between 70 °C and frozen, allowed to warm slowly to room temperature and equilibrate overnight, and were briefly vortexed before analysis. Vitamin D binding protein (DBP) was purchased from Athens Research and Vitamin D Binding protein (VDR) was purchased from Abcam. Proteins were reconstituted according to the manufacturer’s instruction and aliquots of the dissolved protein were added to the hydrated lipid samples to give lipid:protein ratios of the order of 30,000:1.

### NMR Spectra Acquisition, Processing and Analysis

All NMR data were acquired on a 200 MHz, wide bore, Bruker (Karlsruhe, Germany) spectrometer operating at 4.7 T with a ^1^H resonance of 200.1 MHz, a ^13^C resonance of 50.3 MHz and a ^31^P resonance of 81.0 MHz. ^13^C NMR magic angle spinning (MAS) spectra were recorded in a 4 mm two-channel cross polarization-MAS probe with a spinning frequency of 5.5 kHz, using a standard single pulse program with a ^13^C 90° pulse of 2.0 μs at 80 W. Power-gated ^1^H decoupling was applied at 10 W. A recycle delay of 2.0 s was used, 8192 scans were acquired for all spectra, and line broadening of 5 Hz was applied. ^13^C spectra chemical shifts were internally calibrated to the acyl methyl carbon resonance (note that an alternative internal calibration to the choline C_γ_ instead did not alter any of the observations reported here). Single-pulse ^31^P spectra were recorded in the same probe (under static conditions), with a 90° pulse of 2.0 μs at 24.6 W. The recycle delay was again 2.0 s, 2048 scans were acquired and line broadening of either 50 or 75 Hz was applied. All data were analyzed using Bruker Topspin software v4.1.1 (Karlsruhe, Germany). ^31^P chemical shift anisotropy (CSA) parameters were extracted from fits performed with single nuclei (i.e. no overlapping CSAs) in the SOLA NMR plugin within Topspin.

### X-Ray Acquisition

All data were acquired at the X6A beam-line at NSLS at the Brookhaven National Laboratory (Long Island, NY, US). Data were recorded with a 350 mm detector distance, slit size of 130×130 mm and an energy of 9 keV. For each sample, 3 images were acquired in bin mode, with the oscillation range of 0.05°. Data were extracted from 2D using Fit2D (freely available software from ESRF) and converted to *d*-spacing using the standard expression.

### X-Ray Electron Density Profiles

Electron density plots were obtained using the approach of Harper *et al*.^4^, and refined according to the method proposed by Rappolt^5^ and following Pabst *et al*^11^. Appropriate lipid structural parameters were taken from Greenwood *et al*.^12^ with reference to Nagle *et al*.^13^.

## Results and Discussion

Figure 1 displays raw X-ray diffraction (XRD) data for a variety of samples, temperature values and cholesterol concentration values. Two significant features of all of the plotted XRD data (Figure 1) are that each show: (a) broadly lamellar structure with d-spacing in the range ∼60-80 Å and repeat spacings at 1/2 and 1/4; and (b) inhomogeneity in the sense that the first order peaks are both broad and often appear qualitatively to consist of two or more overlapping peaks. We are not aware of an exactly comparable system, but the d-spacing range observed here is consistent with previous reports for related lipid mixtures (e.g. close to C16:0-SM/Chol mixtures ^14^, and somewhat higher than 1,2-dioleoyl-*sn*-glycero-3-phosphocholine (DOPC)/DPPC/Chol mixtures or DPPC/Chol mixtures ^15^. Equally, the presence of substantial inhomogeneity is not unexpected. Although DPPC and SM have been reported to be miscible, the addition of Chol, particularly in ternary systems, is known to induce phase separation ^14,15^, and previous DSC studies of DPPC/SM/Chol systems (albeit with lower concentrations of SM and Chol) have also reported evidence of microdomain formation ^4^. In other ternary systems, phase coexistence below T_m_ in the range of Chol concentrations studied here is usually identified as lamellar gel (L_β_) plus liquid-ordered (L_o_) coexistence ^1^.

**Figure 1:**
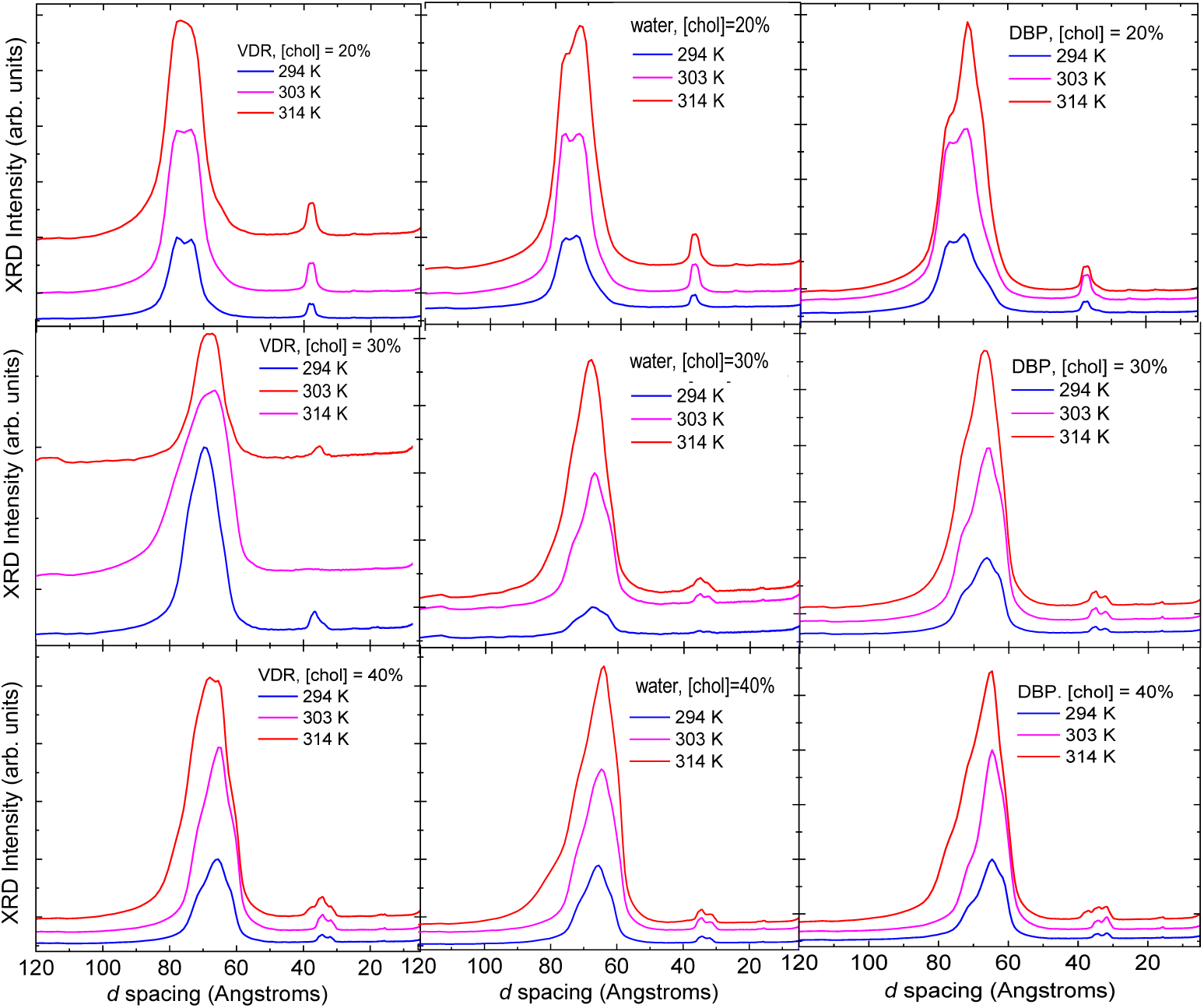
X-ray diffraction scans for all samples, cholesterol concentrations and temperatures in this study. These are raw data and are plotted in “waterfall” mode for clarity.

However, phase separation in ternary combinations of DPPC or SM with 5-cholesten-3-one and ceramide or Chol and 1,2-dipalmitoyl-*sn*-glycerol was identified as coexistence between an L_β_ and an intermediate phase with properties between those of the L_β_ and the L_o_ phases ^16^. Interestingly, in the same work, no evidence of phase coexistence was observed in systems containing either DPPC or SM with equimolar quantities of Chol and ceramide below T_m_.

In Fig. 2. we show XRD data for the purpose of examining evidence for/against presence of crystalline cholesterol and to assess lateral chain packing in the WAXS region. Crystalline Chol (usually in its monohydrate form) has characteristic reflections at ∼34 Å ^17^ as well as relatively sharp reflections in the wide-angle region (2 – 4 Å), which can be observed in coexistence with lamellar phase peaks if present in the sample (see Figure 2) ^18^. In the data presented here, there is no evidence of crystalline Chol in the 20 mol% Chol samples, and there is an overlap between the second order lamellar peaks at 30 and 40 mol% Chol and the region where Chol crystal reflections would appear.

**Figure 2:**
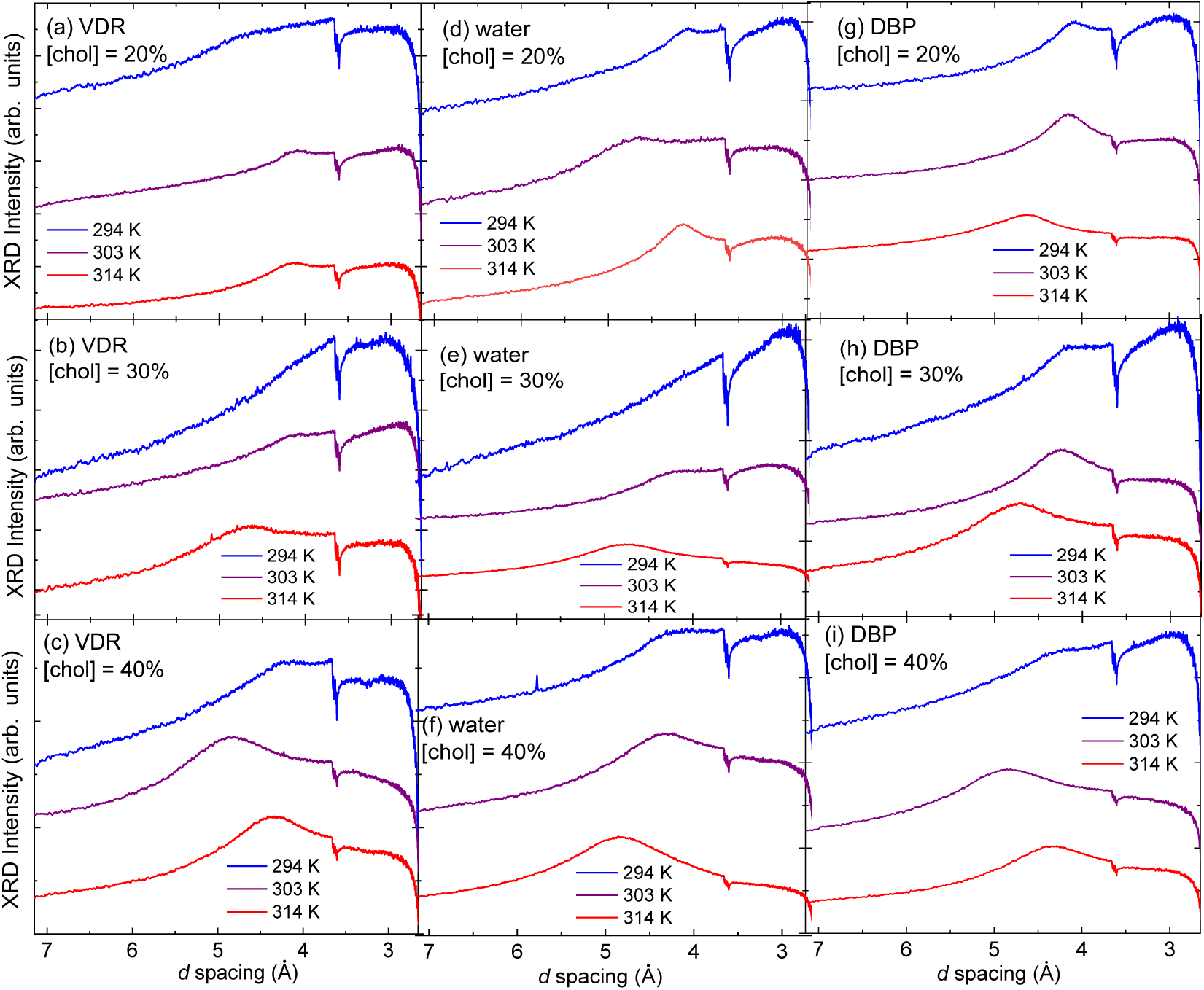
X-ray diffraction scans focusing on the wide-angle region (2.5 to 7 Å) for all samples, cholesterol concentrations and temperatures in this study. These are raw extracts and are plotted in “waterfall” mode for clarity.

At the same time, there is no evidence of crystalline Chol in the wide-angle region in any sample, although the region below ∼3.0 Å is towards the limit of instrumental detection in our set-up. For most samples, the wide-angle region of the diffraction pattern shows a single broad, asymmetric peak between 4 and 5 Å that is characteristic of lateral acyl chain packing. Where the wide-angle peak can be resolved for the 20 and 30 mol% Chol samples at the two lower temperatures, this peak is centred between 4.1 and 4.2 Å, shifting to 4.6 – 4.9 Å at 314 K. For the 30 mol%, the peaks at 294 K are centred at 4.3 Å, shifting first to 4.4 – 4.5 Å at 303 K and then to 4.9 Å at 314 K. Similar dependence of the peak position corresponding to acyl chain lateral spacing on both cholesterol and temperature has been observed in DPPC/Chol systems, although there significantly higher temperatures and cholesterol content were required to achieve a lateral spacing of 4.6 Å ^19^ and the increased spacing may be a result of the unsaturated acyl chain components of bSM. It is notable that there is no overlap of peaks in the wide-angle region, and that all peaks are broad. These observations suggest that in all conditions studied, any coexisting lamellar phases present exhibit substantial but similar acyl chain lateral packing – likely induced in part by Chol and in part by the unsaturated components and varying chain lengths of the bSM – and that the disorder of the chains increases over the 16 K temperature range from values associated with the L_β_ phase to values that are associated with the extent of disorder present in the L_o_ or even L_α_ phases ^20^.

The heterogeneous nature of the samples we look at, coupled with the practical difficulty of taking large number of XRD scans with a sufficiently small spot size for X-ray photons, contributes to the less-than-ideal XRD line-shapes.

^31^P NMR spectra of the 30 and 40 mol% Chol samples with 66 wt% water at 294, 304 and 314 K confirm the presence of a lamellar phase based on the characteristic appearance of the CSA (Figure 3). Additional spectra for samples in the presence of DBP or VDR are available upon request. The anisotropic chemical shift tensor, Δσ, relates to the width of the asymmetric peak and reports on the phosphate group environment ^21^. The values of Δσ range from 40-48 ppm in all but one sample (30% Chol in the presence of DBP), for which poor signal to noise led to an inferior fit. These values are consistent with values reported previously for DPPC/Chol ^22^ and for SM/Chol ^23^ systems over comparable ranges of temperature and cholesterol content. The extracted values of η, which is the asymmetry parameter of the chemical shift tensor, are in the range 0.23 to 0.30. Given the single-nuclei fitting performed, and the results of x-ray diffraction on the same samples, it is likely that the non-zero values of η are linked to overlapping CSAs from two lamellar phases with similar values of Δσ and/or the presence of an L_β_ phase ^24^.

**Figure 3.**
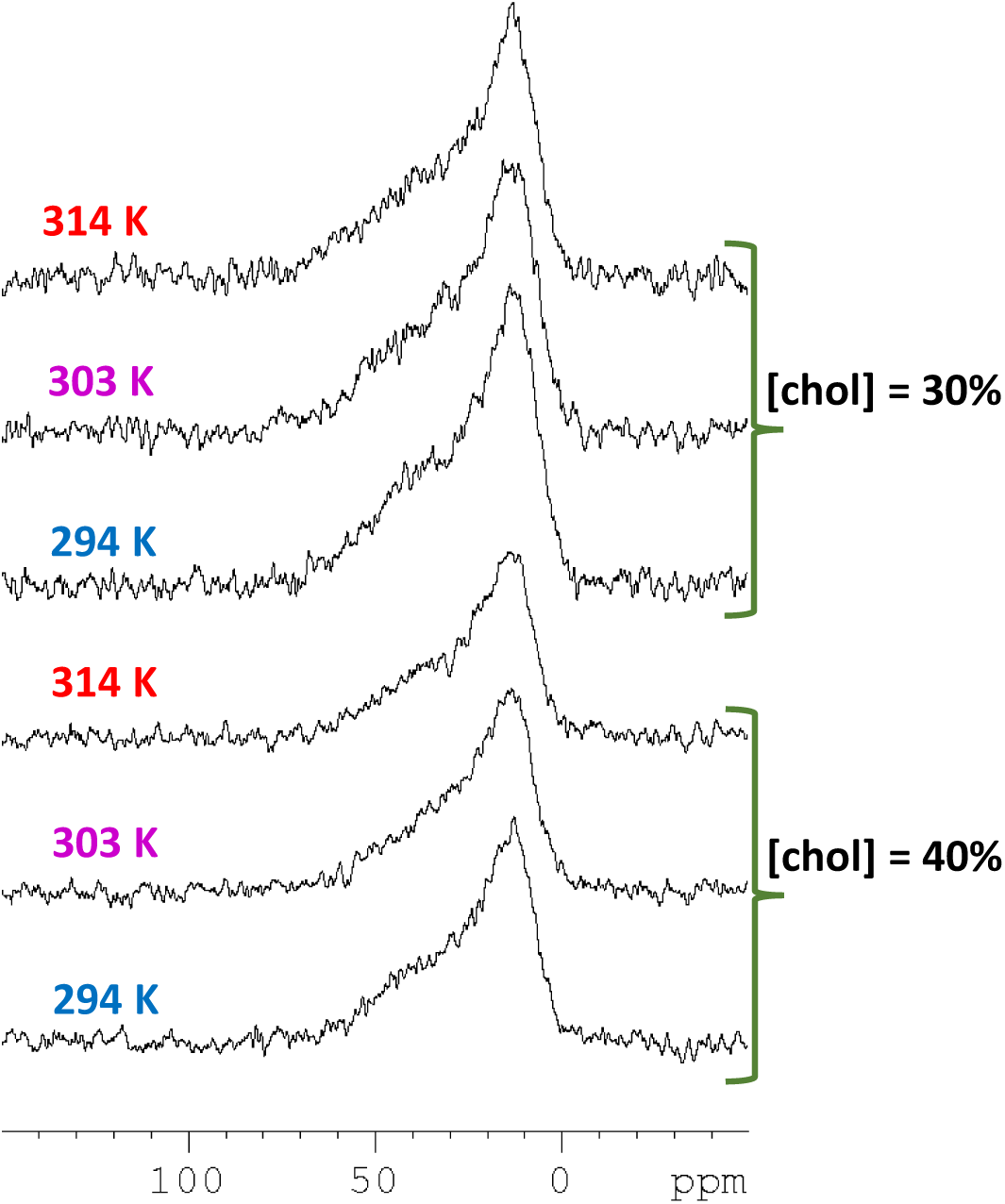
^31^P static NMR spectra of samples containing equimolar quantities of DPPC and bSM with 30 or 40 mol% Chol in 66 wt% water.

The ^13^C NMR MAS spectra for samples containing 30 or 40 mol% Chol at 298, 303 or 314 K with 66 wt% water are shown in Figure 4. Additional spectra for samples in the presence of DBP or VDR are available upon request. Peaks arising from the choline head-group and interfacial carbon atoms are visible in the region 50-73 ppm, with the sharpest peaks arising from the choline C_γ_ (*N*-methyl) at 54.3 ppm, the choline C_α_ at ∼59.5 ppm and the choline C_β_ at ∼66.3 ppm (see ^25^ for choline carbon labelling).

**Figure 4.**
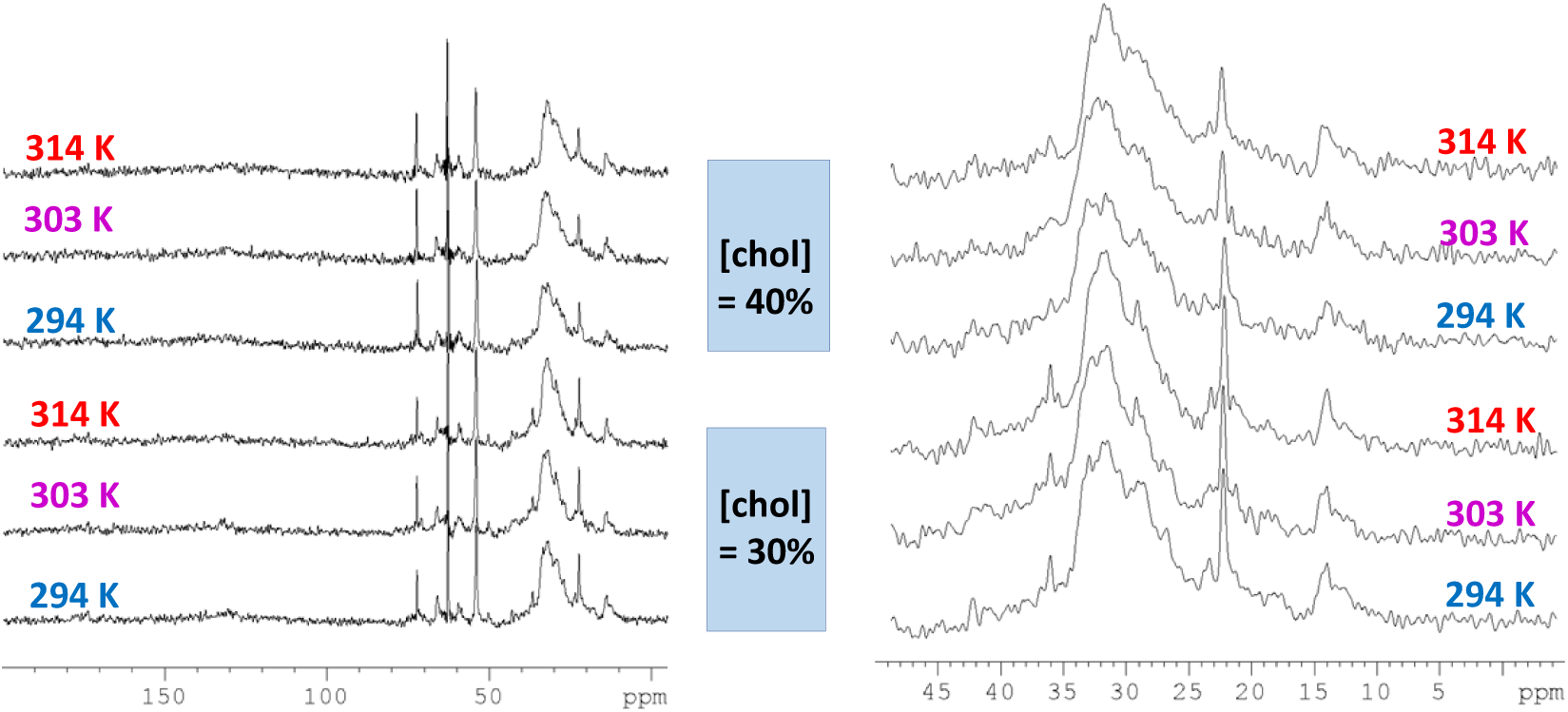
^13^C MAS NMR spectra of samples containing equimolar quantities of DPPC and bSM with 30 or 40 mol% Chol in 66 wt% water. Left panel shows full chemical shift range and right panel shows expanded methylene peak region.

The broad and complex peaks centred at ∼32 ppm result from the acyl chain methylene carbon atoms, probably with some contribution from Chol, and the sharper peak at 22 ppm can be assigned to acyl methylene carbon atoms close to the terminal methyl with a likely contribution from the C26 and C27 Chol methyl groups (see ^26^ for Chol carbon numbering).

The peak at 14 ppm (internally calibrated) can be ascribed to the terminal methyl groups of the lipid acyl chains. This methyl peak tends to be asymmetric (individual methyl groups from different lipid acyl chains are not resolved) and could also include a small contribution from the Chol C18 methyl carbon expected at ∼ 12 ppm. Peak assignments are made with reference to reported chemical shifts for Chol ^26^, SM with Chol ^27^, POPC with Chol ^28^, and other saturated diacylglycerol-phosphatidylcholines ^25^.

There are some notable absences in the reported peaks. In particular, the phospholipid acyl/amide carbonyl resonances expected at between 170 and 180 ppm are not well-resolved in any spectrum, and the alkene resonances expected for sphingosine, cholesterol and any unsaturated component of bSM in the region 120-140ppm are not visible, although there is some distortion of the baseline in the 120-135 ppm range. Combined with the broadness and lack of systematic, temperature-dependent change in chemical shift of the bulk methylene peak ^19,29^, these data suggest/confirm that the lipid interfacial (e.g. carbonyl) and acyl chain carbon atoms in these conditions experience motional restriction that is more similar to that associated with an L_β_ gel phase than with a more fluid L_α_ or L_o_ phase (e.g. see ^27^ for a comparison of SM spectra above and below T_m_).

The x-ray scattering data were further analysed to extract approximations for the electron density profiles in the conditions studied. Due to the data quality (number of reflections, peak broadness and inhomogeneity), an iterative process was employed (see Figure 5) and an example of the output of the analysis is shown in Figure 6. In order to generate the electron density profiles shown, one additional form factor was estimated beyond the measured number of reflections (so a total of 3 or 4 reflections were used for each profile). With these constraints, the estimated head-group positions seemed the most appropriate output, although these should be interpreted cautiously, as an indication of the trends and the ranges of values, rather than as exact values.

**Figure 5.**
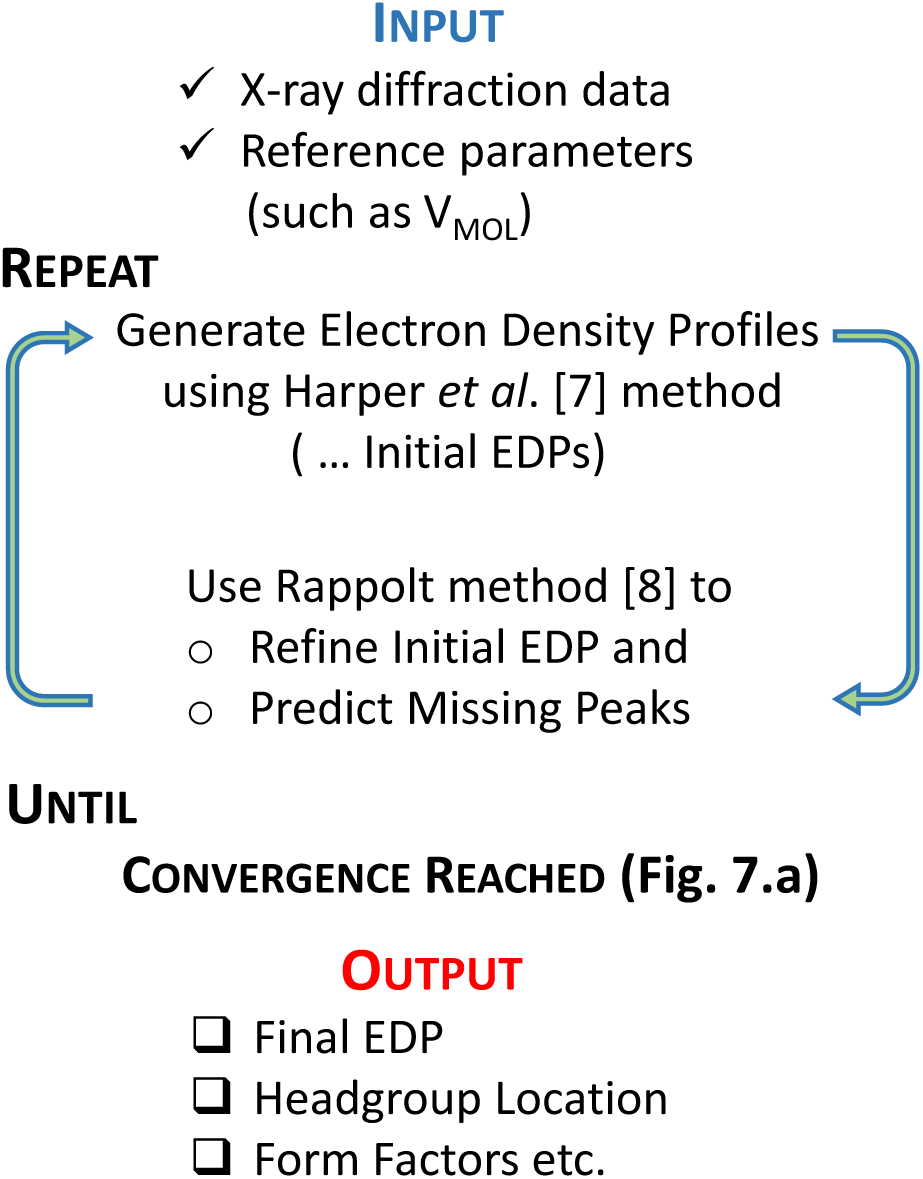
Pseudocode describing the self-consistent X-ray diffraction analysis process, where initial data and known molecular parameters ^12^ are fed into electron density profile calculation, the output of which is then fed to form factor analysis. The process is iterated until a convergence in the position of head-groups is recorded.

**Figure 6.**
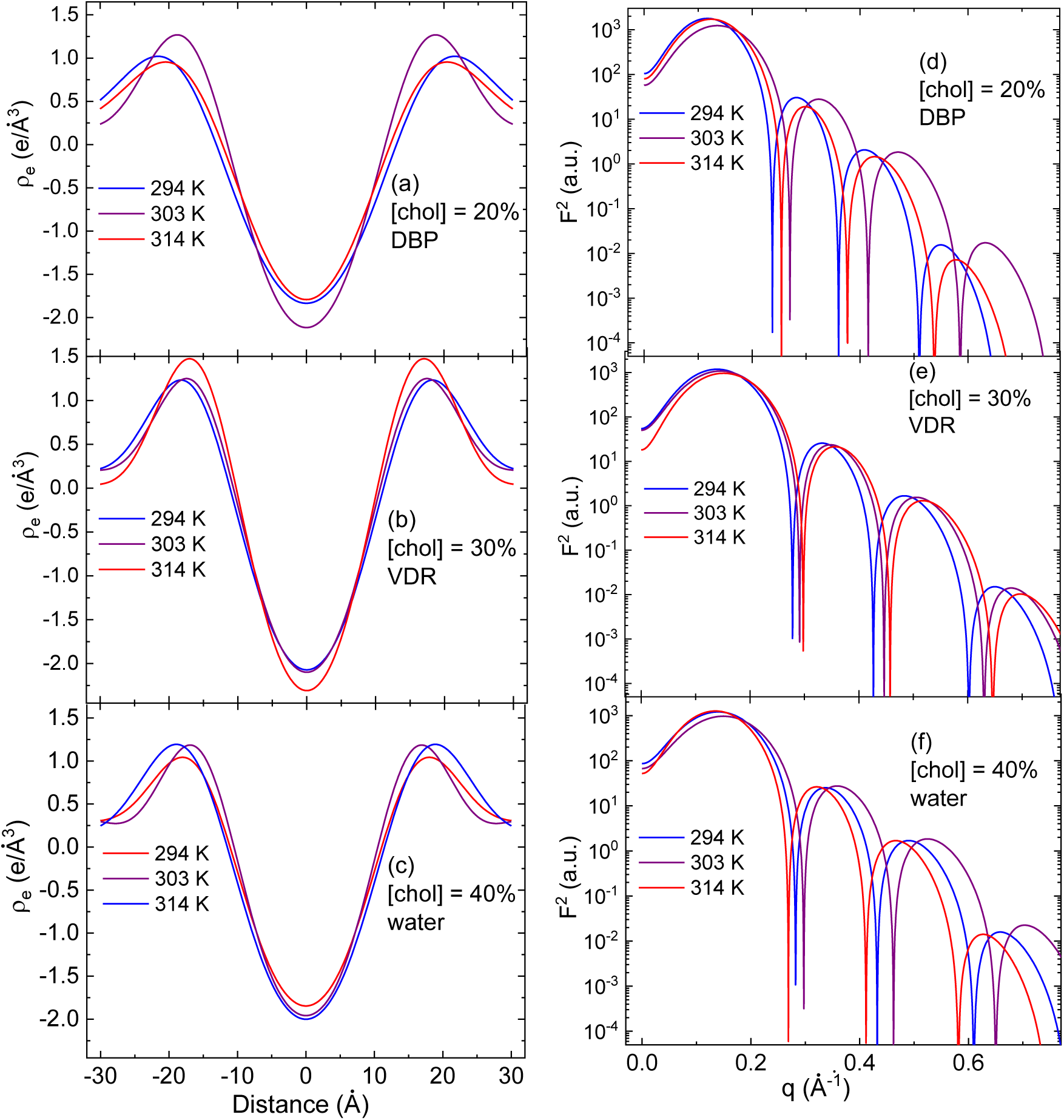
(a) – (c) Electron density profiles obtained based on initial raw data input and Harper et al.^7^ approach for DBP, VDR and water at cholesterol concentrations of 20%, 30% and 40%, respectively. Each panel contains density profiles at three temperature values; (d) – (f) Square of the form factor obtained following the approach by Rappolt ^8^, for the same samples and experimental conditions.

**Figure 7.**
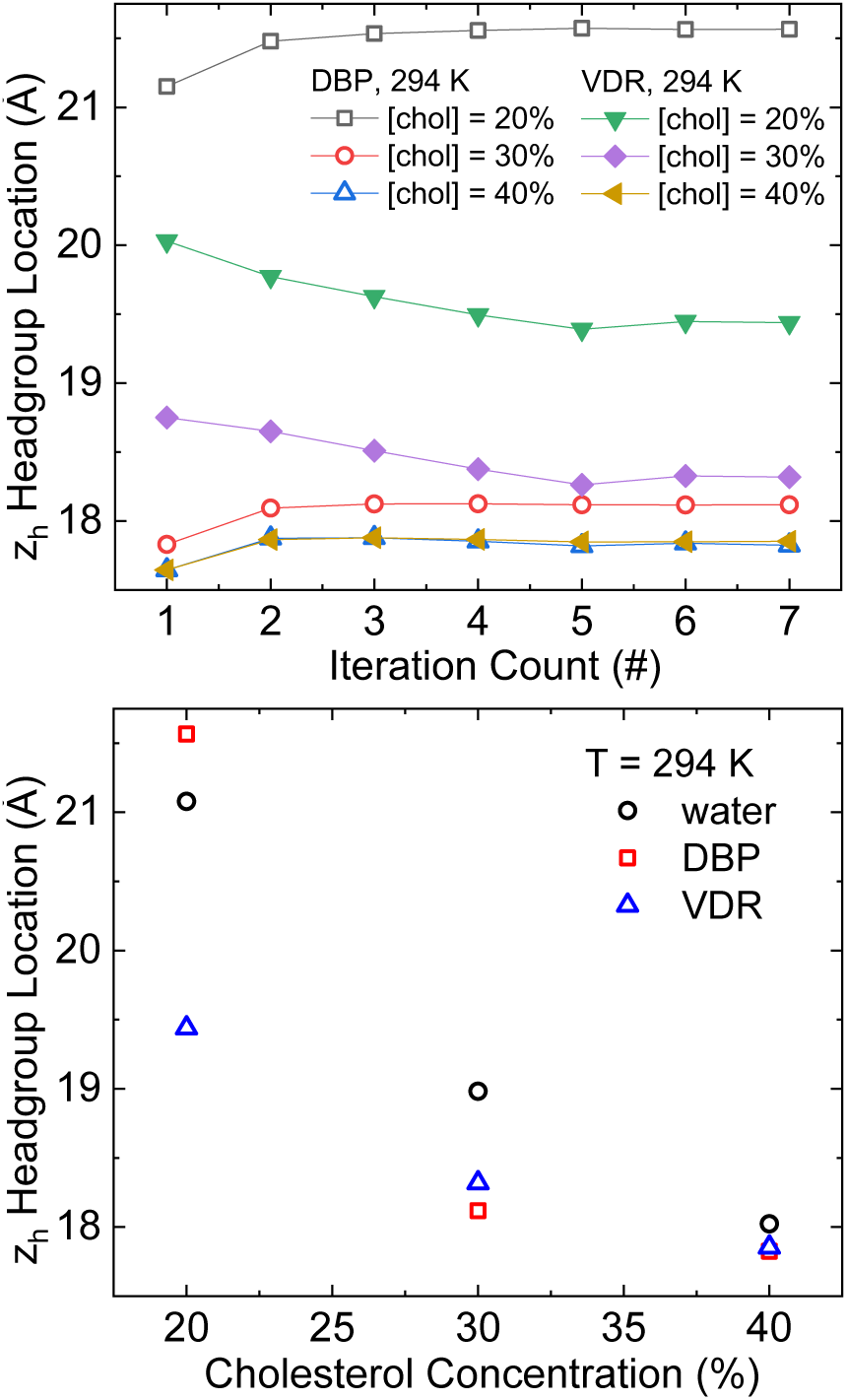
(top) variation of the head-group locations across VDR and DBP samples for varied temperature and cholesterol concentrations. The uncertainty in all points is below 0.5 Angstroms. In most cases there are no significant variations after 5^th^ iteration of the self-consistent approach discussed above, but, for the sake of thoroughness, we have checked samples up to 10^th^ iteration. (lower) the changes in the head-group location as a function of the cholesterol concentrations for three samples at room temperature.

Given the structure and the role these samples have, it is of interest to check the possible changes of the head-group position, as a function of the externally controlled parameters, such as temperature and cholesterol concentration. For convenience, we have organized the analysis so that the successive iterations between electron density profiles as well as formfactors stop when the relative change in the position of the head-group for a given sample and the set of experimental conditions doesn’t exceed 1 part in 10^4^. We note that the results are the same even with more stringent requirement.

For the sake of clarity, we plotted only the last iteration for each case of experimental parameters in Fig. 6, in order to show similarity with the previous result by Rappolt^8^. Additional data are available that demonstrate the convergence.

We focused the analysis of the model output on the variations in the headgroup locations. We observe that for DBP at 294 K, the head-group location changes by 17% over the course of cholesterol concentration change from 20% to 40%. For VDR, at the same temperature, the change is about 8%. There is less variation between samples at 40% Chol than at 30 or 20% Chol, supporting the possibility that the addition of Chol reduces heterogeneity in this system, as has been observed in other systems.

As expected, the general trend in headgroup position is for bilayer thickness to decrease as cholesterol content increases. It is notable that there is more difference between output for samples in the three different conditions (water, DBP or VDR) at 20% Chol than at 30 or 40% Chol. As with a qualitative inspection of the x-ray scattering peaks, it is apparent that the samples displayed more heterogeneity at lower Chol content.

## Conclusions

XRD and NMR were used to investigate the structure of the DPPC:bSM:Chol ternary mixture of phospholipids, with the controlled parameters such as: (a) Vitamin D relevant proteins (DBP and VDR); (b) physiological temperatures between 294 and 314 K; and (c) Chol concentrations between 20 and 40 % by mol. wt. In general – and with some exceptions – increasing temperature tended to reduce the appearance of distinct shoulders in the small-angle region of the ray pattern, and to result in larger and broader wide-angle peaks relating to acyl chain packing. Increasing Chol content had a similar effect, particularly at 40 mol%. These results are reflected in the general decrease in headgroup position with increasing Chol content, reflecting increased chain disorder and a concomitant reduction in bilayer thickness. As might be expected given the low concentrations used, there was no significant effect of the vitamin D relevant proteins.

### Prospective views and future work

It is not clear to us why there is no change in ^13^C methylene peak position, while the XRD suggests measurable change in bilayer thickness and chain packing. It is possible additional work on gel phase only would help clarify this.

Given the relatively rich variations in the structural details, we plan to analyse the same samples with X-ray beams significantly better focused (spot size of the order of 100 nm, as opposed to 2-4 *μ*m range used in the present study). This would likely help us resolve the size and the dynamics of likely domain structure.

Studies in a broader range of temperatures and possibly those done by differential scanning thermometry in addition to XRD and NMR would help elucidate details of the ternary phase diagram.

## Abbreviations

CSA: Chemical shift anisotropy
DBP: Vitamin D binding protein
DOPC: 1,2-Dioleoyl-*sn*-glycero-3-phosphocholine
DPPC: 1,2-Dipalmitoyl-*sn*-glycero-3-phosphocholine
DSC: Differential scanning calorimetry
L_α_: Lamellar fluid disordered phase
L_β_: Lamellar gel phase
L_o_: Lamellar liquid-ordered phase
LUV: Large unilamellar vesicle
MAS: Magic angle spinning
MLV: Multilamellar vesicle
NMR: Nuclear magnetic resonance
SM: Sphingomyelin
T_m_: Phospholipid gel-fluid melting transition temperature
VDR: Vitamin D receptor
XRD: X-ray diffraction

## Acknowledgements

Brookhaven National Laboratory is supported by US Department of Energy (DoE), Office of Science User Facility operated for the DoE by Brookhaven National Laboratory under Contract SC0012704. DLG and AFI acknowledge funding from Khalifa University 2014-KUIRF-L1 grant. We thank Prof. R. V. Law from Imperial College London for generous access to labs and NMR facilities. We also thank S. Rashid and Y. Abdel-Raouf for technical assistance in the early stages of this project. KIH and AFI acknowledge support from Colgate University. AFI acknowledges hospitality of BNL and Cornell University for part of the time worked on this project.

## Data Availability and Supporting Information

Data from this manuscript are available upon reasonable request.

## References

1. Marsh, D. Cholesterol-induced fluid membrane domains: A compendium of lipid-raft ternary phase diagrams. Biochimica et Biophysica Acta - Biomembranes vol. 1788 Preprint at https://doi.org/10.1016/j.bbamem.2009.08.004 (2009).

2. Levental, I., Levental, K. R. & Heberle, F. A. Lipid Rafts: Controversies Resolved, Mysteries Remain. Trends in Cell Biology vol. 30 Preprint at https://doi.org/10.1016/j.tcb.2020.01.009 (2020).

3. García-Arribas, A. B., Alonso, A. & Goñi, F. M. Cholesterol interactions with ceramide and sphingomyelin. Chemistry and Physics of Lipids 199, (2016).

4. Pérez-Isidoro, R. & Costas, M. The effect of neuroleptic drugs on DPPC/sphingomyelin/cholesterol membranes. Chemistry and Physics of Lipids 229, (2020).

5. Hendrich, A. B., Michalak, K. & Wesolowska, O. Phase separation is induced by phenothiazine derivatives in phospholipid/sphingomyelin/cholesterol mixtures containing low levels of cholesterol and sphingomyelin. Biophysical Chemistry 130, (2007).

6. Vist, M. R. & Davis, J. H. Phase Equilibria of Cholesterol/Dipalmitoylphosphatidylcholine Mixtures: 2H Nuclear Magnetic Resonance and Differential Scanning Calorimetry. Biochemistry 29, (1990).

7. Harper, P. E., Mannock, D. A., Lewis, R. N. A. H., McElhaney, R. N. & Gruner, S. M. X-ray diffraction structures of some phosphatidylethanolamine lamellar and inverted hexagonal phases. Biophysical Journal 81, (2001).

8. Rappolt, M. Bilayer thickness estimations with “poor” diffraction data. Journal of Applied Physics 107, (2010).

9. Bartoccini, E. et al. Nuclear lipid microdomains regulate nuclear vitamin D 3 uptake and influence embryonic hippocampal cell differentiation. Molecular Biology of the Cell 22, (2011).

10. Borel, P., Caillaud, D. & Cano, N. J. Vitamin D Bioavailability: State of the Art. Critical Reviews in Food Science and Nutrition 55, (2015).

11. Pabst, G. et al. Structural analysis of weakly ordered membrane stacks. Journal of Applied Crystallography 36, (2003).

12. Greenwood, A. I., Tristram-Nagle, S. & Nagle, J. F. Partial molecular volumes of lipids and cholesterol. Chemistry and Physics of Lipids 143, (2006).

13. F. Nagle J. et al. Revisiting Volumes of Lipid Components in Bilayers. The Journal of Physical Chemistry B 123, 2697–2709 (2019).

14. Maulik, P. R. & Shipley, G. G. N-palmitoyl sphingomyelin bilayers: Structure and interactions with cholesterol and dipalmitoylphosphatidylcholine. Biochemistry 35, (1996).

15. Chen, L., Yu, Z. & Quinn, P. J. The partition of cholesterol between ordered and fluid bilayers of phosphatidylcholine: A synchrotron X-ray diffraction study. Biochimica et Biophysica Acta - Biomembranes 1768, (2007).

16. Busto, J. v. et al. Lamellar gel (Lβ) phases of ternary lipid composition containing ceramide and cholesterol. Biophysical Journal 106, (2014).

17. Shieh, H. S., Hoard, L. G. & Nordman, C. E. The structure of cholesterol. Acta Crystallographica Section B Structural Crystallography and Crystal Chemistry 37, (1981).

18. Gater, D. L., Seddon, J. M. & Law, R. V. Formation of the liquid-ordered phase in fully hydrated mixtures of cholesterol and lysopalmitoylphosphatidylcholine. Soft Matter 4, (2008).

19. Clarke, J. A., Heron, A. J., Seddon, J. M. & Law, R. v. The diversity of the liquid ordered (Lo) phase of phosphatidylcholine/cholesterol membranes: A variable temperature multinuclear solid-state NMR and x-ray diffraction study. Biophysical Journal 90, (2006).

20. Mills, T. T. et al. Order parameters and areas in fluid-phase oriented lipid membranes using wide angle x-ray scattering. Biophysical Journal 95, (2008).

21. Seelig, J. 31P nuclear magnetic resonance and the head group structure of phospholipids in membranes. BBA - Reviews on Biomembranes vol. 515 Preprint at https://doi.org/10.1016/0304-4157(78)90001-1 (1978).

22. Guo, W. & Hamilton, J. A. A Multinuclear Solid-State NMR Study of Phospholipid-Cholesterol Interactions. Dipalmitoylphosphatidylcholine-Cholesterol Binary System. Biochemistry 34, (1995).

23. Costello, A. L. & Alam, T. M. Investigating the impact of cholesterol on magnetically aligned sphingomyelin/cholesterol multilamellar vesicles using static 31P NMR. Chemistry and Physics of Lipids 163, (2010).

24. Holland, G. P., McIntyre, S. K. & Alam, T. M. Distinguishing individual lipid headgroup mobility and phase transitions in raft-forming lipid mixtures with 31P MAS NMR. Biophysical Journal 90, (2006).

25. Leftin, A. & Brown, M. F. An NMR database for simulations of membrane dynamics. Biochimica et Biophysica Acta - Biomembranes vol. 1808 Preprint at https://doi.org/10.1016/j.bbamem.2010.11.027 (2011).

26. Gater, D. L. et al. Hydrogen bonding of cholesterol in the lipidic cubic phase. Langmuir29, (2013).

27. Guo, W. et al. A solid-state NMR study of phospholipid-cholesterol interactions: Sphingomyelin-cholesterol binary systems. Biophysical Journal 83, (2002).

28. Epand, R. M., Bain, A. D., Sayer, B. G., Bach, D. & Wachtel, E. Properties of mixtures of cholesterol with phosphatidylcholine or with phosphatidylserine studied by 13C magic angle spinning nuclear magnetic resonance. Biophysical Journal 83, (2002).

29. Ishikawa, S. & Ando, I. Structural studies of dimyristoylphosphatidylcholine and distearoylphosphatidylcholine in the crystalline and liquid-crystalline states by variable-temperature solid-state high-resolution 13C NMR spectroscopy. Journal of Molecular Structure 271, (1992).

